# Neuronal modulation in the mouse superior colliculus during covert visual selective attention

**DOI:** 10.1101/2021.02.05.429996

**Authors:** Lupeng Wang, James P. Herman, Richard J. Krauzlis

**Affiliations:** Laboratory of Sensorimotor Research, National Eye Institute Bethesda, Maryland 20892 USA

## Abstract

Covert visual attention is accomplished by a cascade of mechanisms distributed across multiple brain regions. Recent studies in primates suggest a parcellation in which visual cortex is associated with enhanced representations of relevant stimuli, whereas subcortical circuits are associated with selection of visual targets and suppression of distractors. Here we identified how neuronal activity in the superior colliculus (SC) of head-fixed mice is modulated during covert visual attention. We found that spatial cues modulated both firing rate and spike-count correlations, and that the cue-related modulation in firing rate was due to enhancement of activity at the cued spatial location rather than suppression at the uncued location. This modulation improved the neuronal discriminability of visual-change-evoked activity between contralateral and ipsilateral SC neurons. Together, our findings indicate that neurons in the mouse SC contribute to covert visual selective attention by biasing processing in favor of locations expected to contain relevant information.

## Introduction

Visual selective attention is the ability to selectively process relevant stimuli and ignore irrelevant distractors, and is achieved through a distributed network of brain areas that includes both cortical and subcortical areas. Our understanding of the neuronal mechanisms of visual selective attention so far mainly comes from neurophysiological studies in non-human primates that manipulate attention using informative cues. For instance, cortical neurons, including those in the visual cortex^1^, frontal^2^ and parietal cortices^3^, display cue-related modulation of their visual responses during selective attention tasks. The main features of cue-related modulation in cortical neurons include increases in firing rate^4^ in favor of the cued stimulus and changes in the correlated variability of neuronal ensembles^5^, both of which can affect the decoding of visual information. Subcortical areas are also involved in selective attention, either in parallel or in conjunction with cortical mechanisms. Neurons in primate superior colliculus^6^, thalamus^7,8^ and caudate nucleus of the basal ganglia^9^ also display cue-related modulation during visual selective attention tasks. In contrast to cortical mechanisms that appear to regulate the quality of local visual processing, subcortical circuits have been implicated in the spatial weighting of visual signals^10^ and the suppression of distractors^11,12^ during perceptual tasks.

The mouse has emerged as a promising model for studying the neuronal mechanisms of visual selective attention, because the genetic tools available in mice provide unparalleled opportunities to study selective attention at molecular, genetic, cellular and circuit levels. Complementing these tools, several behavioral studies over the last few years have demonstrated the feasibility of studying visual selective attention in mice, both in head-fixed^13,14^ and freely moving^15^ preparations. Notably, we recently reported that mice display perceptual benefits from informative spatial cues – a well-known attentional effect – in three different visual spatial attention tasks typically used in primates^13^. These efforts pave the way for further investigating the neuronal mechanisms of visual selective attention in mice using experimental approaches that are not yet readily available in primates. For instance, optogenetic manipulation of the basal ganglia “direct pathway” in mice during spatial attention tasks has revealed a circuit that biases visual processing in favor of the cued visual location^16^. In the visual cortex of mice, results from a visual attention task have identified subthreshold membrane dynamics that depend on the spatial-cue context^14^.

The midbrain superior colliculus is a crucial subcortical structure for the control of visual selective attention in several species^10,17^ but the role of the mouse SC in selective attention has not yet been established. Since the SC has direct or indirect connections with all known brain areas involved in attention, understanding what and how the SC contributes to visual selective attention could be a linchpin for understanding the overall circuit mechanisms. On the other hand, studies of mouse SC visual functions have largely focused on visuomotor processing related to innate visual behaviors, such as predator avoidance or prey approach^18^. We recently reported that inhibiting visually evoked SC activity in mice impairs their voluntary visual perceptual choices, and that the perceptual impairment was larger when a competing visual stimulus was present^19^, consistent with a role of the mouse SC in visual selection^20^. However, the involvement of the mouse SC in visual selective attention itself has not yet been explicitly tested. It is possible that the mouse SC is involved in prioritizing the representation of expected stimuli, or suppressing the distractors; alternatively, the mouse SC might simply be involved in the early processing of visual events. To distinguish among these hypotheses, it is crucial to investigate how SC neuronal activity is modulated during visual selective attention tasks.

Here we investigated the neuronal correlates of visual selective attention in the mouse SC by recording the spiking activity of neurons during a visual orientation change-detection task and using spatial cues to manipulate the allocation of selective attention. We found that visually evoked activity in the mouse SC displayed cue-related modulation, including changes in spike rate and interneuronal spike-count correlations. By comparing activity across attention task conditions, we determined that the cue-related modulation was the result of enhancement at the cued spatial location rather than suppression at the uncued location. Together, our results demonstrate that neurons in the mouse SC are involved in visual selective attention, and that SC neurons can contribute to attention by biasing signal processing in favor of spatial locations expected to contain behaviorally relevant events.

## Results

To investigate cue-related modulation of mouse SC neuronal activity, we recorded the activity of SC neurons in two variants of a spatial cueing task. The main task, which we term “contra/ipsi cue”, was similar to one we used previously^13,21^. In brief, head-fixed mice viewed stimuli on a pair of lateralized displays while running on a wheel. The animals’ locomotion controlled each trial’s progression through several epochs defined by visual stimulus events (Fig. 1a). Presentation of a single lateralized Gabor patch served as a spatial cue, indicating the potential location of an upcoming orientation change, and defined the start of the “cue epoch”. The appearance of a second Gabor patch in the opposite visual hemifield marked the start of the “2-patch epoch”, throughout which both Gabor patches remained present. In trials with an orientation change (50% of trials), the start of the “change epoch” was marked by a tilt in orientation of the cue patch. Mice were required to lick a center spout within a 500 ms response window to indicate their detection of the orientation change and receive a liquid reward. Each session was organized into alternating sub-blocks of 40 left-cue and 40 right-cue trials. We used this version of the task to characterize the spatial specificity and time course of cue-related modulation in mouse SC neurons (n = 94 sessions). In a subset of sessions (n = 25) we also recorded SC neuronal activity in a variant of the main task that included sub-blocks of 80 no-cue trials interleaved with left-cue and right-cue sub-blocks; accordingly, we refer to this variant as the “cue/no-cue” task.

**Fig. 1.**
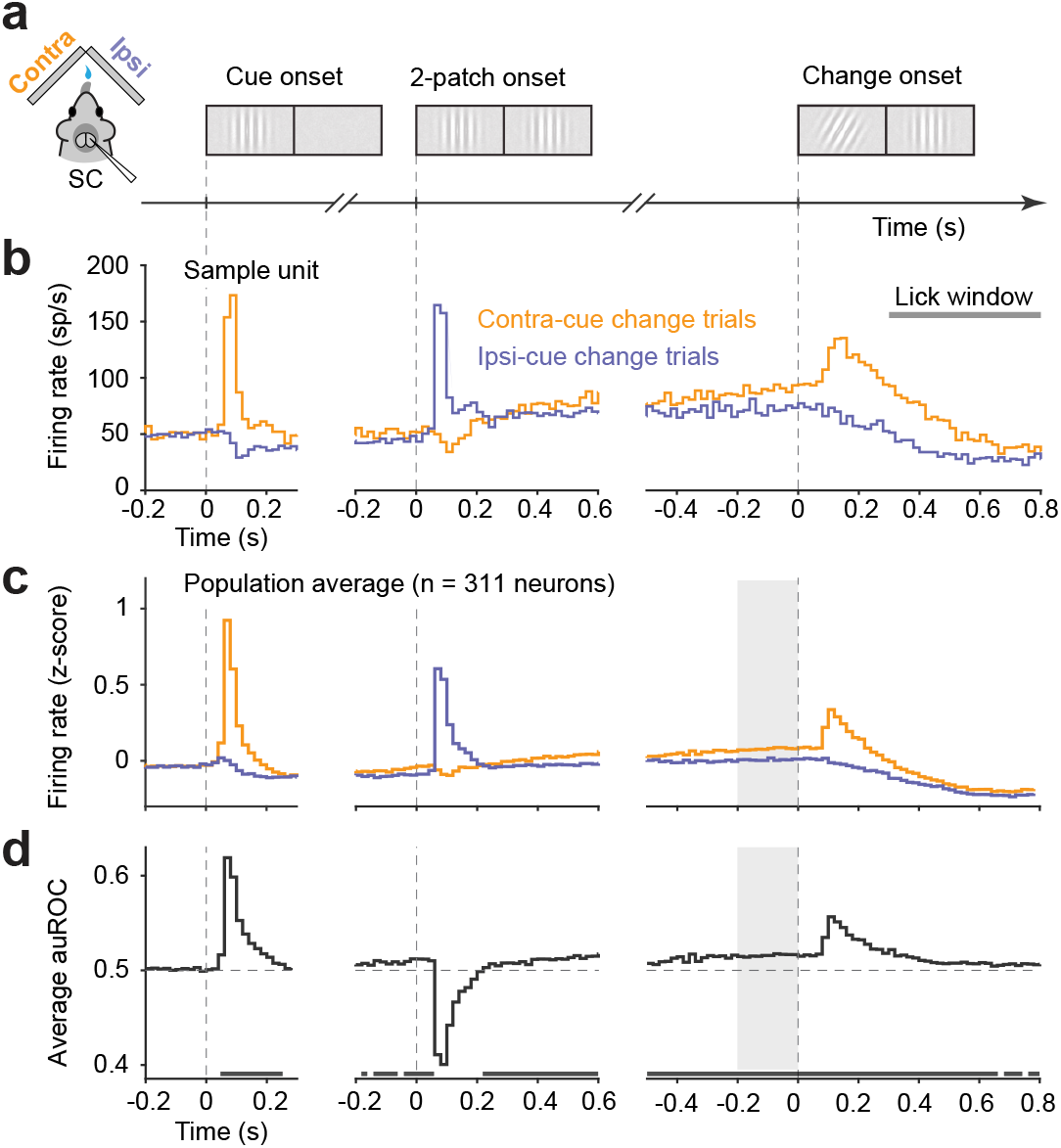
Spatial cue related modulation of mouse SC neuronal activity in a visual change detection task. **a**) Schematics of unilateral recording in the mouse SC during the “contra/ipsi cue” orientation change detection task, and illustration of visual stimuli on two visual displays in sequences of task epochs: cue, 2 patch and change. **b**) Firing rates of a sample SC unit aligned on three task epochs, shown as Peristimulus time histograms (PSTHs) of contra-cue change trials (orange) and ipsi-cue change trials (blue) in 20-ms bins. Gray horizontal bar indicates lick window from 300 ms to 800 ms after orientation change. **c**) Normalized population PSTHs aligned on the onset of three epochs. Plotting conventions as in **b**. Gray area indicates the final 200 ms of the 2-patch epoch (defined as the delay period). **d**) Time course of population average area under ROC (auROC) comparing spike counts between contra-cue change and ipsi-cue change trials aligned on three task epochs in 20-ms bins. The row of gray boxes below mark bins in which population auROC are significantly > 0.5 chance level, as measured in one-tailed signed rank tests.

Extracellular activity was recorded from SC neurons located at least 400 μm below the dorsal surface of SC with moveable chronic 16-channel microwire bundles. At these depths, recorded neurons were located in the intermediate and deep layers of SC. For the purpose of documenting neuronal activity related to the visual attention task, we analyzed the activity of 311 / 481 SC neurons with clear visual spatial receptive fields that overlapped the location of the contralateral visual stimulus (Fig. S1).

### SC neuronal responses to visual events and modulation by spatial cue location

In addition to exhibiting phasic activity for stimulus-onset and orientation-change events, many SC neurons displayed cue-related modulation during multiple epochs of the attention task. The results from a sample unit illustrate the pattern observed across the population of SC neurons during the “contra/ipsi cue” task (Fig. 1b). Prior to cue onset, many neurons displayed tonic activity (population mean spike count within a 200 ms interval before cue onset: 2.39 ± 0.33, mean ± 95% confidence interval [CI]), and most neurons (79%, 247/311, see Methods) were not modulated by the location of the spatial cue (Fig. 1c), even though the neurons were recorded during sub-blocks of 40 consecutive trials with the same cue location. The onset of the spatial cue caused a phasic increase of activity in most neurons (81%, 251/311, significant units, see Methods) when it was presented contralaterally (contra-cue), and a small decrease when presented ipsilaterally (ipsi-cue). The onset of the second Gabor patch had a similar effect, causing a phasic increase in activity when presented contralaterally in ipsi-cue trials and a decrease when presented ipsilaterally in contra-cue trials.

Following these onset transients, neuronal activity in contra-cue trials gradually increased, on average exceeding activity in ipsi-cue trials ~250 ms after the start of the 2-patch epoch and remaining elevated throughout the remainder of the task epoch. We identified this elevation of neuronal activity in contra-cue trials compared to ipsi-cue trials during the latter half of the 2-patch epoch as cue-related modulation for two reasons. First, the visual stimuli presented during this interval of the two trials types were identical, so the elevation in activity in the 2-patch epoch depended on the location of the spatial cue presented in the preceding epoch. Second, this cue-related modulation was not a result of lingering visual responses caused by the preceding cue, because it emerged gradually over time during the 2-patch epoch on a time scale that anticipated the possible visual change event.

After the near-threshold change in orientation, many SC neurons (35%, 108 / 311) exhibited robust transient increases in activity for changes in the contralateral visual field and modest slower reductions in activity for changes in the ipsilateral visual field (Fig. 1b,c, right panels). Note that these change-related increases and decreases in SC activity were superimposed on different baseline levels of neuronal activity, because of the differences in cue-related modulation during the preceding 2-patch epoch.

To quantify cue-related modulation, we used the receiver operating characteristic (ROC) approach from signal detection theory^22^. We computed the area under the ROC curve (auROC) by comparing spike rates in ipsi-cue trials (“signal absent”) to spike rates in contra-cue trials (“signal present”) in consecutive non-overlapping 20 ms bins (Fig. 1d). The average auROC across our population of SC neurons indicates that before cue onset, the spike rates of SC neurons did not differentiate between contra-cue and ipsi-cue trial types (p > 0.1 in all bins within 200 ms, one-tailed Wilcoxon Signed rank test for population auROC). Rather, the spike rates on contra-cue trials started to become higher than ipsi-cue trials at 220 ms after the onset of the 2-patch epoch (p < 0.05, one-tailed Wilcoxon Signed rank test on population auROC values) and remained significant throughout the rest of the epoch. These results demonstrate that, even though spatial cue information was available throughout the sub-block of trials, cue-related modulation of SC neurons emerged only after the visual cue stimuli were presented.

Attention-related neuronal modulation is often documented during a “delay-period”, an epoch during attention tasks after the transient effects of stimulus onsets have waned, during which the sustained modulatory effects of attention can be assessed on otherwise tonic neuronal activity. In our experiments, we defined the “delay period” as the final 200 ms of the 2-patch epoch, when mice viewed the two Gabor patches and waited for a potential orientation change to occur. Across our sample of SC units, the attention modulation during the delay period measured by auROC was significantly larger than the chance (0.5) level (p < 10^−15^, one-tailed Wilcoxon Signed rank test, Fig. 2a), indicating that at the population level mouse SC neurons have higher spike rates in contra-cue trials than ipsi-cue trials during this interval (37%, 115/311 of total units show auROC values significantly > 0.5, bootstrapped 95% CI > 0.5). Similarly, the distribution of delay period attentional modulation indices, the other widely used measurement of attention related modulation of neuronal activity, supports the same conclusion (Fig. 2b). These results demonstrate that mouse SC neuronal activity displays a classic hallmark of attention-related modulation.

**Fig. 2.**
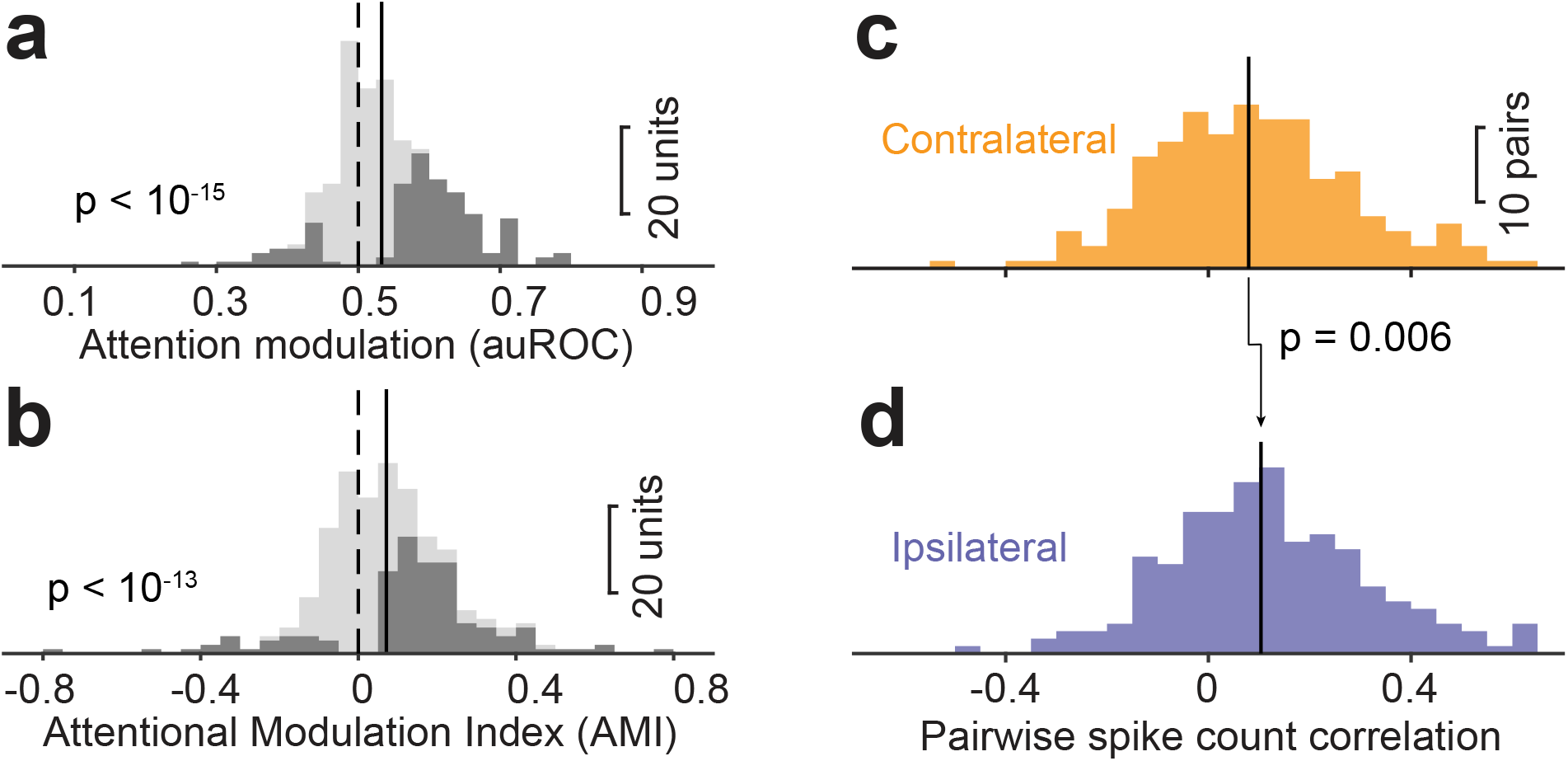
Population summary of cue-related modulation on mouse SC firing rate and interneuronal spike-count correlations. **a**) Distribution of delay period activity auROC comparing contra-cue and ipsi-cue trials; bin width of histograms is 0.05. Dark bars count units with auROC value significant different from chance level (bootstrapped 95% CI ⊄ 0.5). Dashed line indicates chance value of 0.5, solid line indicates population median (0.53). **b**) Distribution of delay-period AMIs between contralaterally and ipsilaterally cued trials, using the same conventions as in **a**. Dark bars count units with significant AMI (p < 0.05, rank sum test of spike count between ipsi-cue and contra-cue trials). Dashed line indicates chance value of 0, solid line indicates population median (0.07). **c**) Distribution of delay period interneuronal spike-count correlations during “contra-cue” trials of simultaneously recorded SC neuronal pairs. Solid line: population median (0.079). **d**) Distribution of delay period spike-count correlations during “ipsi-cue” trials. Solid line: population median (0.104). P value is from paired-sample Wilcoxon signed rank test comparing population medians between “contra-cue” and “ipsi-cue” trials.

In addition to modulation of spike rate, spatial cueing can also influence the information available in neuronal populations through effects on the structure of correlated variability amongst neurons^5,23^. To assess how the correlated variability amongst mouse SC neurons was modulated by spatial cueing, we computed spike count (“noise”) correlations during the delay period of contra-cue trials and ipsi-cue trials in simultaneously recorded neuronal pairs (n = 203 pairs). As shown in Fig. 2c-d, the distribution of spike-count correlations across pairs of mouse SC neurons was broad (standard deviation for contra-cue = 0.196; ipsi-cue = 0.198) and on average positive (mean for contra-cue = 0.084; ipsi-cue = 0.116). Notably, spike-count correlations in contra-cue trials (median = 0.079) were significantly smaller than those in ipsi-cue trials (median = 0.104), indicating that the degree of correlated variability among mouse SC neurons was reduced when mice awaited a potential visual event that might occur in their receptive fields. This result indicates that spatial cueing can potentially alter information available in mouse SC neuronal populations by reducing correlated variability.

Because several other factors might also contribute to the modulation of mouse SC neuronal activity during our attention task, including behavioral states and locomotion^24,25^, we sought to compare the potential influence of these factors, as well as the spatial cue condition, on the spike rates of individual SC neurons using linear regression analysis (Fig. S2). This analysis revealed that some mouse SC neurons were indeed significantly modulated by running speed (29%, 90/311, significant units, see Methods) or pupil size (27%, 83/311). However, the influence of these factors on spike rate was manyfold smaller than that of spatial cue condition, which was the single largest contributor (p <10^−9^ for both Tukey-Kramer post-hoc comparison tests following one-way ANOVA on linear regression coefficients) to variation in spike rate during the delay period. These results reinforce our conclusion that spatial cue information was specifically important in modulating SC neurons during the visual attention task.

### Cue-related modulation results from enhanced activity at the location of expected visual events

The difference in spike rate we observed between contra-cue and ipsi-cue trials during the delay period could be due to enhancement of processing at the spatial location expected to contain behaviorally relevant information, or suppression of activity at locations not expected to contain such information, or a combination of both effects. Distinguishing between these possibilities is important for determining the underlying circuit mechanisms of selective attention operating amongst our SC neurons.

To address this point, we used a “cue/no-cue” variant of our attention task^13^ that allowed us to compare SC neuronal activity evoked by cued and uncued stimuli to activity evoked by identical visual stimuli but presented in a context with no spatial cueing. For the no-cue trials in these experiments, the cue-epoch was replaced with pink noise and then followed by our standard 2-patch epoch (Fig. 3a). Orientation changes in these no-cue trials occurred in pseudorandom order on the left or right side with equal frequency. Left-cue, right-cue, and no-cue trials were organized as interleaved sub-blocks.

**Fig. 3.**
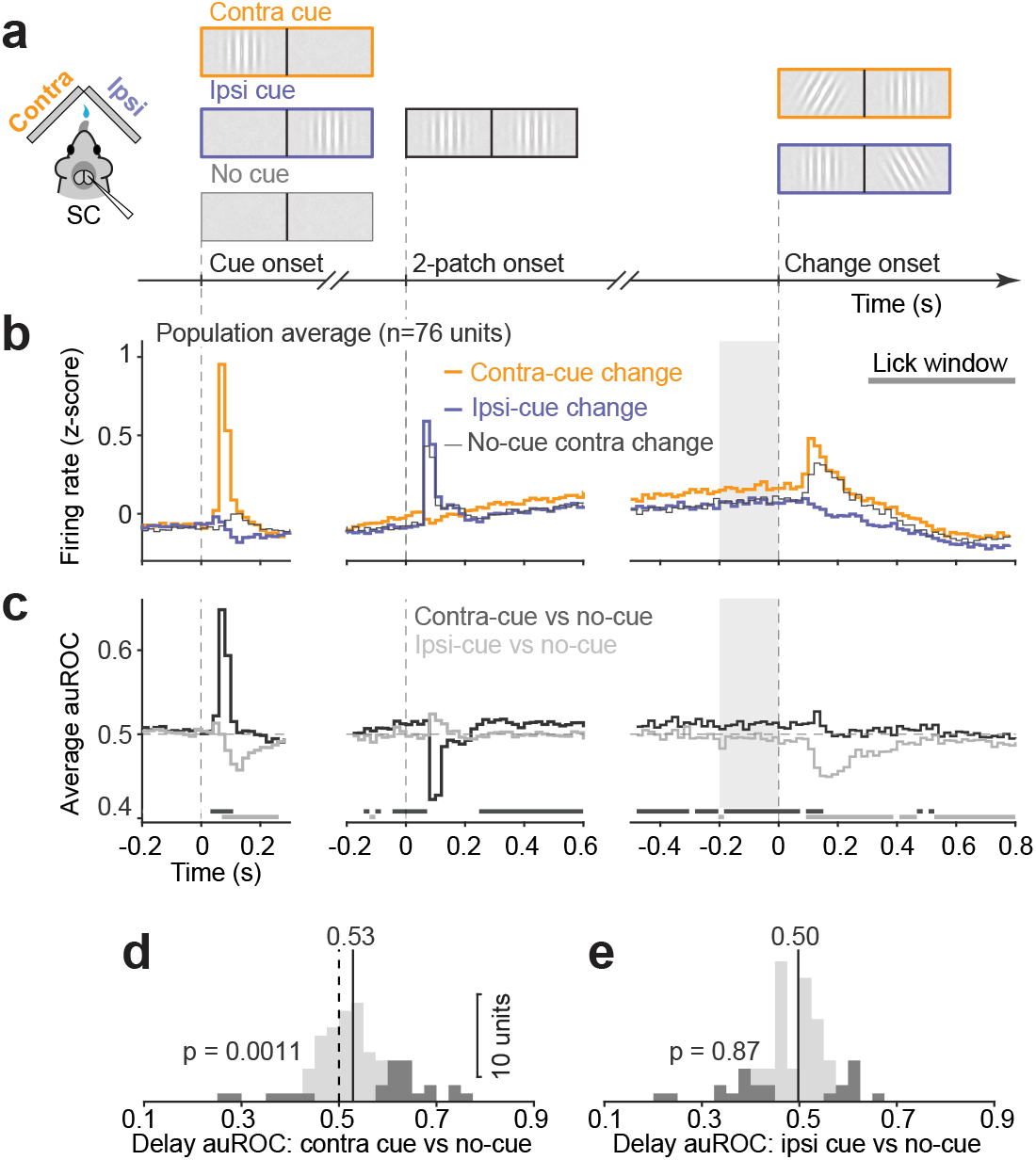
Cue-related modulation resulted from enhanced activity at contralaterally cued spatial locations. **a**) Schematic of epochs in “cue/no-cue” task, where no-cue trials had cue epoch replaced with a noise epoch and orientation change could occur either left or right with equal frequency. No-cue sub-blocks were interleaved with cued sub-blocks. **b**) Normalized population PSTHs of SC neurons for contra-cue change (orange), ipsi-cue change (blue) and no-cue contra change trials (gray), aligned on the onset of task epochs. Gray area indicates the delay period used for analysis shown in **d**-**e**. Gray horizontal bar indicates the 500-ms lick window. **c**) Time course of population average auROCs comparing “contra-cue” spike counts to “no-cue” change trials (dark), and comparing “ipsi-cue” to “no-cue” change trials (gray), aligned on three task epochs in 20 ms bins. Only “no-cue” trials with contralateral orientation change were illustrated. The row of dark gray boxes below mark bins in which population auROCs for contra-cue vs no-cue are significantly > 0.5 chance level, as measured in one-tailed signed rank tests; row of light gray boxes below mark bins in which population auROC for ipsi-cue vs no-cue are significantly < 0.5 chance level. **d**) Distribution of delay period auROCs comparing “contra-cue” and “no-cue” change trials. Dark bars count units with auROC values significantly different from chance level (bootstrapped 95% CI ⊄ 0.5). Dashed line indicates chance value of 0.5, solid line with value indicates population median. **e**) Presentation as in **d**, but for auROCs comparing “ipsi-cue” to “no-cue” trials.

We first verified that providing the spatial cue improved behavioral performance. Perceptual sensitivity, measured using the behavioral metric d’ from signal detection theory, was significantly higher on with-cue trials compared to no-cue trials (with-cue: 1.64 ± 0.11, mean ± SEM; no-cue: 1.36 ± 0.11, p < 0.001, Wilcoxon signed rank test), consistent with our previous results^13^. The decision criterion was also significantly different, shifting in favor of the cued location on trials when spatial cueing was provided (with cue: −0.36 ± 0.11; no-cue: −0.15 ± 0.13, p = 0.004).

At the neuronal level, we found clear evidence that cue-related modulation in the mouse SC was the result of enhancement at the cued location, with little or no suppression at the uncued location. As expected, given the cueing conditions, SC activity in the contra-cue and ipsi-cue trials during the “cue/no-cue” task (Fig. 3b) recapitulated the cue-related modulation found during the “contra/ipsi cue” task (Fig. 1c). The novel finding from this set of experiments was that activity in the no-cue trials (thin gray line in Fig. 3b) was not only lower than the activity in the contra-cue trials, it was nearly identical to the activity in the ipsi-cue trials. The only exceptions were the transient differences after the onset of the cue and the occurrence of the contralateral change, which would be expected given the difference in visual stimulus conditions at those points in the trial.

To document the time course of these cue-related effects, we again computed spike rate auROC values in consecutive 20 ms bins (Fig. 3c), separately comparing contra-cue and ipsi-cue to the no-cue condition. Before the onset of the cue epoch, neither comparison showed average auROC values significantly different from the chance level (p > 0.05 in all bins within −200 ms to 0 ms interval, one-tailed Wilcoxon Signed rank test on population auROC values), consistent with our findings in the “contra/ipsi cue” dataset that cue-related modulation was not present before the start of each trial. The average auROC values for contra-cue versus no-cue became significantly greater than chance at 240 ms after the onset of the 2-patch epoch (p < 0.05, one-tailed Wilcoxon Signed rank test on population auROC values) and remained significantly elevated for the duration of the epoch, indicating a sustained enhancement of activity at the cued location. In contrast, the auROC values for ipsi-cue versus no-cue trials were not different from chance in any time bin after 120 ms during the 2-patch epoch, demonstrating that cueing did not produce suppression at the uncued location.

Finally, we examined delay period neuronal modulation in the “cue/no-cue” task variant. We computed individual SC neuron auROC values during the final 200 ms of the 2-patch epoch (“delay period”), and found the distribution to be significantly greater than chance when comparing contra-cue to no-cue (p=0.0011, one-tailed Wilcoxon Signed rank test, Fig. 3d), but not different from chance when comparing ipsi-cue to no-cue (p = 0.87, Fig. 3e). We found the same pattern of results when we quantified population cueing effects with an attention modulation index (AMI) rather than auROC (contra-cue vs no-cue: 0.036 ± 0.019, p = 0.006, one-tailed Wilcoxon signed rank test; ipsi-cue vs no-cue: −0.004 ± 0.016, p = 0.71).

Together, these results demonstrate that the cue-related modulation observed in mouse SC neurons was due to the enhancement of processing at the cued spatial location rather than suppression at the uncued location.

### Cue-related modulation improves SC neuronal discriminability of visual events

Having established that spatial cues enhanced mouse SC neuronal activity specifically at the cued location, we next examined how cueing influenced neuronal activity evoked by the behaviorally relevant stimulus event. We recently found that unilateral suppression of SC neuronal activity in a short time interval immediately after the visual orientation change caused major deficits in the ability of mice to correctly detect these near-threshold visual events^19^. Given that SC neuronal activity appears to be crucial for this detection task, we sought to identify how the effects of spatial cueing on SC neurons might contribute to the observed improvements in task performance during “cue/no-cue” experiment.

One possibility is that spatial cueing improves the ability of mouse SC neurons to discriminate between change and no-change events at the cued location, as has been observed in primate visual cortex^26^. However, we found no evidence that spatial cueing affected the neuronal discriminability for contralateral events. We computed auROC values for a “change” window (150 ms interval beginning 60 ms after the change, or a matched interval in trials with no change; see Methods) by comparing spike counts in change versus no-change trials, separately for contra-cue and no-cue trials. There was no significant difference between the average auROC values of the contra-cue condition compared to no-cue (p = 0.82, two-tailed Wilcoxon Signed rank test), despite the fact that the spike counts in contra-cue change trials were significantly higher than those in no-cue change trials (p = 0.03, one-tailed Wilcoxon Signed rank test).

Another possibility is that spatial cueing improves the ability of SC neurons to discriminate change events in the contralateral visual field, relative to SC neuronal activity occurring at the same time in the other SC (Fig. 4a), as has been observed in primate SC^27^. To test this, we used unilateral recordings of SC activity obtained separately during contralateral and ipsilateral orientation changes as a proxy for simultaneous bilateral recordings during contralateral change events (Fig. 4b, d). We again computed auROC values for the “change” window, comparing contra-change and ipsi-change spike counts, separately for the cued and no-cue trials. This analysis revealed that auROC values were significantly higher in cued trials compared to no-cue trials (p = 0.0045, Fig. 4c, e). In addition, a larger proportion of SC neurons displayed significant auROC values (bootstrapped 95% CI ⊄ 0.5) in cued trials than in no-cue trials (χ-square test; p = 0.009). Therefore, spatial cueing significantly improved SC neuronal discriminability when the relative levels of activity in the two colliculi were taken into consideration.

**Fig. 4.**
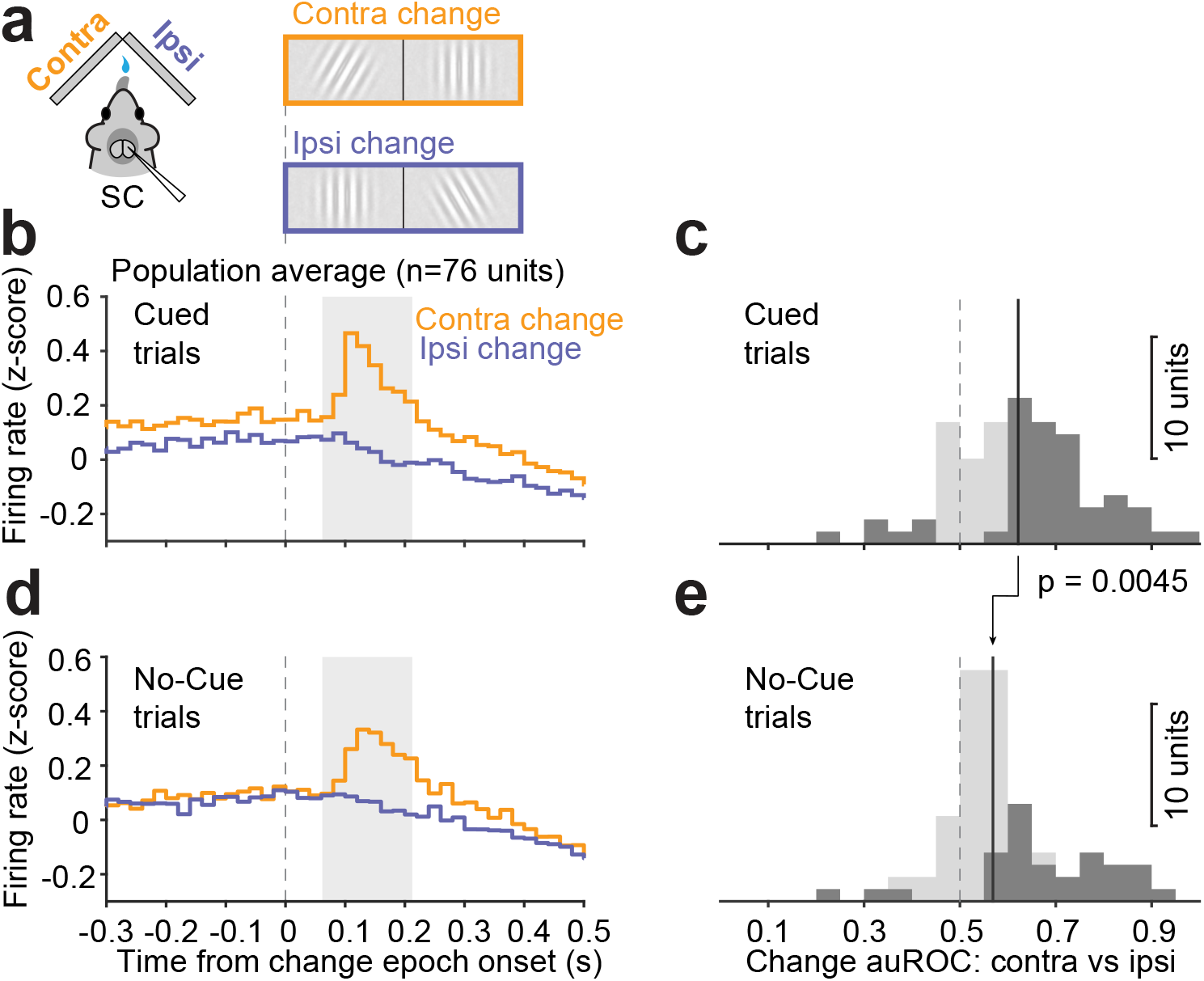
Cue-related modulation enhanced the neuronal detection of the orientation change based on comparing activity between the contralateral and ipsilateral SC. **a**) Schematic of apparatus and events in “cue/no-cue” task. **b**) Average normalized population PSTHs of SC neurons during “contra-cue” change trials (orange) and “ipsi-cue” change trials (blue). Traces are aligned on the change onset. Gray area: 150 ms change interval (60 – 210 ms after change onset) used to compute auROC comparing contralateral and ipsilateral change trials. **c**) Distribution of change auROCs in cued trials. Dark bars are units with auROCs significantly different from chance level. Dashed line indicates chance value of 0.5, solid line indicates population median (0.62). **d**) Presentation as in **b** but comparing “no-cue contra” (orange) to “no-cue ipsi” (blue) change trials. **e**) Presentation as in **c**, but in no-cue trials; (median = 0.57). The p value indicates comparison of population median between **c** and **e**, two-tailed Wilcoxon signed rank test.

Together, these results demonstrate that cue-related modulation in the mouse does not necessarily increase the ability of SC neurons to locally discriminate between change and no-change events, but instead enhances activity at the cued location so that neuronal discriminability is improved when comparing activity across both sides of the SC.

## Discussion

Our study reveals that neurons in the mouse SC display cue-related modulation during a covert visual selective attention task – notably, the perceptual benefits from spatial cueing were confirmed by improvements in detection of a near-threshold visual change. The cue-related modulation emerged after the spatial cue stimulus was presented and persisted through the visual target event, consistent with the time course of attention allocation in other species^28,29^. The main feature of cue-related modulation was an increase in the spike rates of SC neurons when the mouse was cued to attend the contralateral visual field, consistent with the retinotopic representation in the SC^30^. In addition to effects on spike rate, the average spike count correlations between pairs of SC neurons were also lower with contralateral spatial cues. Furthermore, by comparing activity across attentional conditions, we determined that the cue-related modulation was due to enhancement of SC activity at the cued location rather than suppression of SC activity at the uncued location.

Our results identify how neurons in the mouse SC can contribute to the mechanisms of visual selective attention – their spiking activity is selectively elevated with spatial cueing so that the processing of visual information is biased in favor of spatial locations expected to contain relevant information. This biasing of SC activity does not alter the local neuronal discriminability of visual events, and hence would not improve performance if activity from only one side or region of the SC were used to guide detection performance. Instead, this biasing enhances the difference in activity between the cued location and locations represented elsewhere in the SC, thereby improving neuronal discriminability of the relevant event if the readout mechanism involved a bilateral or global comparison of activity across the entire SC, consistent with previous results in the primate^27^.

Stimulus competition is an important aspect of visual selective attention and the SC plays a crucial role in selecting target stimuli amongst competing distractors in many species^17,20,31^, presumably reflecting an evolutionarily conserved midbrain function^32,33^. In the primate, SC activity is modulated in a variety of paradigms that involve stimulus selection, including the selection of targets for orienting movements^34,35^ and also selection of visual stimuli in the absence of orienting movements^6,36^. This modulation is an indicator of the biased competition between the alternative stimuli that unfolds before the subject’s response and that is usually settled in favor of the cued stimulus. Further evidence of this competition has been provided by causal manipulations of SC activity which, by artificially increasing or decreasing SC for one of the stimulus locations, can cause major changes in which alternative the subject selects^12,37,38^. The fact that we found comparable enhancement of SC spiking activity in favor of the cued stimulus in our covert attention tasks suggests that similar competitive mechanisms also apply to the mouse SC.

However, the particular competitive mechanism that would explain our results is somewhat unexpected. There is compelling evidence that inhibitory feedback circuits to the SC (or optic tectum) play a key role in implementing stimulus selection, and these involve strong suppression of non-cued locations across the SC map^17,39^. Our pattern of results indicates that a different mechanism is responsible for the cue-related modulation of our SC neurons. By interleaving cue and no-cue trials, we determined that our cue-related modulation was due to enhancement of visual processing at the cued location, rather than suppression at the uncued distractor location. This enhancement of activity at the cued location – with no change at the uncued location – is not easily explained by broad inhibitory feedback but would be consistent with a more focused excitatory or disinhibitory circuit mechanism. For example, we recently demonstrated that activity through the direct pathway of the basal ganglia is linked to the allocation of spatial attention^16^; this might involve a disinhibitory mechanism from the substantia nigra to the SC that would be consistent with our current neuronal data. It is also possible that other attention task designs, such as those that require actively ignoring visual stimuli at uncued locations^13^, would reveal evidence for broad inhibition like that found in previous studies.

The logic of the attention mechanism in the SC indicated by our results is different from that described in visual cortex, because it involves a global comparison rather than a local improvement in discriminability. In visual cortex, it is thought that a local improvement of neuronal discriminability through sharpened visual tuning^40^, changes in receptive field properties^41^, and alterations in the statistical structure of activity amongst the population of active neurons^5^ all contribute to the perceptual improvements at cued locations during the allocation of attention^26,42^. The lack of cue-related improvements in local neuronal discriminability of mouse SC neurons in our results might be related to how their activity is used in the task. Neurons in the SC are generally not selective to visual features or exhibit much broader tuning than visual cortical neurons^43^. Thus, the tuning of local pools of SC neurons for visual features might be less relevant for determining the accuracy of perceptual decisions; instead, the relative magnitude of event-related activity across the SC retinotopic map might be a much more important factor in setting the limits of detection performance^44^. This interpretation is consistent with previous studies showing that relative activity across the SC (or optic tectum) plays a central role in visual selection in fish^45^, birds^46^, and primates^27^. This type of mechanism is also reminiscent of computational models that use differential weighting of sensory evidence to explain how spatial cueing can account for the perceptual improvements during selective attention^47^. Thus, the relative activity across the SC might be a central component of the mechanism that implements visual spatial attention.

The effect of spatial cueing on interneuronal “noise” correlations we observed in mouse SC is consistent with previous observations in primate sensory cortical areas, but likely carries different functional implications. Pairs of neurons in primate visual cortex often display smaller correlations for cued stimuli than for uncued stimuli^42,48^, consistent with specific hypotheses about how the correlation structure of neuronal activity in these visual cortical areas impacts the decoding mechanism used to guide behavior^49,50^. Outside of cortical sensory areas, the possible importance of the neuronal correlation structure is less well established but also necessarily depends on how the neuronal activity is decoded^51,52^. Our hypothesis, that global decoding of SC neurons supports the detection of behaviorally relevant events, implies a specific relationship between cue-related modulation of correlations in neuron pairs within and between the two halves of SC^27^. Future visual attention experiments using simultaneous bilateral recordings in the mouse SC might help resolve these issues.

How SC attentional control interacts with cortical mechanisms of attention remains to be explored. Besides prominent roles of the SC in early visual processing in mice^53^, outputs from SC could significantly modulate visual cortical activity via visual thalamus in mice^54–57^. Thus, it is possible that cue-related modulation in the SC may contribute to cortical correlates of visual attention in mice. On the other hand, how cue-related modulation influences mouse visual cortical processing remains unclear. Notably, a recent study found spatial-context-dependent enhancement of visual processing in the mouse primary visual cortex, using a block-organized visual detection task with target stimuli that occurred at one of two locations within the same hemifield^14^. However, no competing distractor stimulus was presented in that study, and it is evident that attention-related modulation in the primate visual cortex is influenced heavily by stimulus competition^58,59^. It is unclear how mouse visual cortical activity would be modulated in attention tasks that include competing visual stimuli, such as ours.

In addition to interacting with visual cortical mechanisms, there is also evidence that the SC can contribute to visual attention through mechanisms that operate further downstream. In the macaque, SC inactivation causes major deficits in visual attention performance but without altering the well-known correlates of attention in visual cortex^60^, indicating that the SC makes its contributions to attention through other cortical or subcortical regions. The basal ganglia were proposed as a possible candidate, based on their role in learned associations between visual events and behavioral responses^61^, and this has been supported by more recent findings. For instance, inactivation of the primate SC disrupts attention-related modulation of neuronal activity in caudate nucleus of the striatum^9,62^, together with causing behavioral deficits in spatial attention. In mice, it has been demonstrated that the dorsomedial striatum, putative homolog of the primate caudate nucleus, is causally involved in visual perceptual choices and visual selective attention^16,21^.

Our knowledge about neuronal mechanisms of visual selective attention in mice remains far from complete. Nevertheless, our findings demonstrate that the mouse SC neuronal activity displays a classic hallmark of attention-related modulation observed in the primate brain, illustrating the utility of the mouse as a model system for dissecting the neuronal mechanisms of attention. Future studies investigating how the SC, basal ganglia and cortical circuits interact could provide important insights into how a higher-order function like selective attention is implemented through the cooperative activity across these diverse brain regions and circuits.

## Material and Methods

### Animals

All procedures were conducted on wild-type C57BL/6J mice (JAX stock # 000664). The mice were housed in a 12:12 reversed day-night cycle, with lights off at 9 am, and all experimental procedures and behavioral training were done in the lights-off portion of the cycle (9am-9pm). Two male and two female mice weighing 18-25 grams were surgically implanted at age 6-8 weeks and then used in experiments for up to ~9 months. All the mice were in group housing (2-4 cage mates) prior to the surgical procedure, and subsequently singly housed after the implant surgery. All experimental procedures and animal husbandry were approved by the NIH Institutional Animal Care and Use Committee (IACUC) and complied with Public Health Service policy on the humane care and use of laboratory animals.

### Stereotaxic surgery

Each mouse was implanted with a head-holder before behavioral training; the procedure was similar to that in our previous studies^13^. During the surgery, animals were anesthetized with isoflurane (4% induction, 0.8-1.5% maintenance) and secured by a stereotaxic frame with ear bars (Kopf Instruments, CA). Dexamethasone (1.6 mg/kg) was administered to reduce inflammation. A feedback-controlled heating pad (PhysioSuite, Kent Scientific, CT) was used to maintain the body temperature at 37°C, and artificial tears were applied to the eyes to prevent them from drying. After the animal’s head was leveled in the stereotaxic frame, a scalp incision was made along the midline. A custom-designed titanium head post for head-fixing was positioned and secured to the skull using Metabond (Parkell Inc., NY). The skin wound edge was then closed with sutures or tissue adhesive (Vetbond, 3M, MN). After surgery, mice received subcutaneous ketoprofen (1.85mg/kg) daily for up to three days to alleviate any potential discomfort.

After each mouse was trained on the detection task for 20-30 days, a second surgery for implanting microwire bundles was carried out. Anesthesia procedure was identical to the first head-post surgery. After the animal’s head was leveled in the stereotaxic frame, a small craniotomy was made for implanting with moveable custom 16-wire microwire bundles (Innovative Neurophysiology, NC). The coordinates for the tips of stainless steel cannula of the microwire bundles were ±0.8~1.1 mm from midline (M-L axis), −3.65~-4.0 mm from Bregma (A-P axis) and 0.2-0.5 mm ventral (D-V axis), based on a standard mouse brain atlas^63^. The cranial opening was sealed with bone wax and the microwire bundle assembly was secured on the skull with Metabond. Mice again received post-surgery subcutaneous ketoprofen (1.85mg/kg) as needed to alleviate potential discomfort.

### Food control

After mice recovered from surgery and returned to above 95% of their pre-surgery weight (typically within 7-9 days), they were placed on a food control schedule. Mice had free access to water, but their intake of dry food was controlled, and they were allowed to augment their dietary intake by access to a nutritionally complete 8% soy-based infant formula (Similac, Abbott, IL). Overall food intake was regulated to maintain at least 85% of their free-feeding body weight, and the health status of each mouse was monitored daily throughout the study. Mice were initially acclimatized to handling procedures by having their heads gently restrained while receiving the soy-based fluid under manual control via a sipper tube. After the initial exposure to soy-based fluid, the animal was more securely head-fixed, and manual delivery was continued. Once mice were adapted to these procedures, we switched to automatic delivery of fluid under computer control in the behavioral apparatus.

### Behavioral apparatus

The behavioral apparatus consisted of a custom-built booth that displayed visual stimuli to the mouse, the updating of the display was coupled to their locomotion. Details of apparatus construction are described elsewhere^64^. The mouse was head-fixed in the center of the apparatus, positioned atop a polystyrene foam wheel (20-cm diameter) that allowed natural walking or running movements along a linear path. An optical encoder (Kübler, Germany) was used to measure the rotation of the wheel. The front walls of the booth incorporated a pair of LCD displays (VG2439, ViewSonic, CA) positioned at 45° angles from the animal’s midline such that each display was centered on either the right or left eye and subtended ~90° horizontal by ~55° vertical of the visual hemifield, at a viewing distance of 27.5 cm. The interior of the booth was lined with sound absorbing material to reduce acoustic noise. The entire apparatus rested on a vibration isolation air table (Newport, CA). The experiments were controlled by a computer using a modified version of the PLDAPS system^65^. Our system omitted the Plexon device, but included a Datapixx peripheral (Vpixx Technologies, Canada) and the Psychophysics Toolbox extensions^66,67^ for Matlab (The Mathworks, MA), controlled by Matlab-based routines run on a Mac Pro (Apple Inc, CA). The Datapixx device provided autonomous timing control of analog and digital inputs and outputs and synchronized the display of visual stimuli. A reward delivery spout was positioned near the snout of the mouse; lick contacts with the spout were detected by a piezo sensor (Mide Technology Co., MA) and custom electronics. Each reward was a small volume (5-10 μl) of an 8% solution of soy-based infant formula (Similac, Abbott, IL) delivered by a peristaltic pump (Harvard Apparatus, MA) under computer and Datapixx control. The temperature inside the apparatus was maintained in a temperature range of 70-80° F.

### Visual detection tasks with spatial cueing

The tasks were similar to those we used previously^13,64^. Experiments were organized in blocks of randomly shuffled, interleaved trials, and each trial consisted of a sequence of epochs that the mouse progressed through by walking or running forwards on the wheel. Each epoch was defined by the particular stimuli presented on the visual displays, and the duration of each epoch was determined by the time required for the mouse to travel a randomized distance on the wheel. A typical trial lasted several seconds.

Each trial followed a standard sequence of four epochs. The average luminance across each visual display in all epochs was 4-8 cd/m^2^. In the first epoch (“noise”, not shown), the uniform gray of the inter-trial interval was replaced by pink noise with an RMS contrast of 3.3%; this epoch was presented for a wheel distance of 10 - 20 cm (range of time: 0.2 - 0.3 s). In the second epoch (“cue”), on cued trials a vertically oriented Gabor patch was added to the pink noise, centered in either the left or right visual display. The Gabor patch consisted of a sinusoidal grating (95% Michelson contrast) with a spatial frequency of 0.1 cycles per degree, a value chosen based on the visual spatial acuity of mice, modulated by a Gaussian envelope with full width at half-maximum of 18° (σ = 7.5°). The phase of the grating was not fixed, but throughout the trial was incremented in proportion to the wheel rotation with every monitor refresh, so that the sinusoidal pattern was translated within the patch by approximately the same distance that the mouse traveled on the wheel; the Gabor patch on the left (right) drifted leftward (rightward), consistent with optic flow during locomotion. This second epoch lasted for 46-92 cm (0.36-1.55 s). On no-cue trials, no Gabor was added during the second epoch, but the otherwise the timing was the same. In the third epoch (“2 patch”), a second Gabor patch with the same spatial frequency and orientation appeared on the other side of the visual display; this epoch lasted for 107 - 214 cm (0.84 - 3.6 s). On no-cue trials, both Gabor patches appeared simultaneously in 2-patch epoch. The visual stimuli in the fourth epoch (“change”) depended on whether or not the trial included an orientation change. If the trial did include an orientation change, the cued Gabor patch changed orientation at the onset of the visual-event epoch; in no-cue trials, either one of the two Gabor patches changed its orientation with equal frequency. The amplitude of the orientation change was always 9°, which was near the detection threshold of mice. If the trial did not contain an orientation change, the two Gabor patches did not change their orientation, so that the “change” epoch unfolded as a seamless extension of the previous 2-patch epoch. Thus, in every experiment, the cue was always 100% valid, but an equal number of change and no-change trials were interleaved, making the probability of a change on any given trial 50% from the perspective of the subject.

The task of the mouse was to lick the spout when he or she detected a change in the orientation of the Gabor patch and to otherwise withhold from licking. Mice were required to lick within a 500-ms response window starting 300 ms after the orientation change in order to score a “hit” and receive a fluid reward. Any lick before the response window would result in trial abort and timeout penalty. If the mouse failed to lick within the response window after an orientation change, the trial was scored as a “miss” and no reward was given but no other penalty was applied. On “no change” trials, if the mouse licked within the same response window aligned on the unmarked transition to the fourth epoch, the trial was scored as a “false alarm”, which led to timeouts; if they correctly withheld from licking throughout the entire “change” epoch, the trial was scored as a “correct reject”. At the end of correct reject trials, the trial was extended to include an additional “safety-net epoch” in which the cued Gabor patch underwent a supra-threshold (30°) orientation change and the mouse could receive a reward by licking within a comparable response window. Responses in the safety-net epoch were not used for any analysis in the study.

Both variants of spatial cueing task experiments were organized as blocks of trials, with the sub-block conditions defined based on our recording site in the SC. In the “contra/ipsi cue” task variant, each block contained 80 trials, subdivided into 40 contralateral-cue trials (i.e., contralateral to our recording site in the SC), 40 ipsilateral-cue trials. The 40 contralateral-cue and 40 ipsilateral-cue trials were run back-to-back, with the order of the two sub-blocks randomly determined at the beginning of each single session. In the “cue/no-cue” task variant, each block contained 160 trials, subdivided into 40 contralateral-cue trials, 40 ipsilateral-cue trials, and 80 no-cue trials. The 40 contralateral-cue and 40 ipsilateral-cue trials were also run back-to-back, with the order of the two sub-blocks randomly determined in a given session. The sub-block of 80 no-cue trials were run either before or after the 80 (i.e., 40 plus 40) cue trials with equal probability. Of all trial types, 50% were with orientation change, randomly interleaved with no change trials. During a daily session, mice typically completed 320 - 800 trials in total.

### Electrophysiological recording

Spiking activity of SC neurons was recorded in four C57BL/6J mice (2 males, 2 females) implanted with moveable 16-wire microwire bundles (Innovative Neurophysiology, NC). Electrophysiological signals were acquired through an RZ5D processor and Synapse Suite interface (Tucker-Davis Technologies, FL) with voltages band-pass filtered (0.3 to 7 kHz) and sampled at 25k Hz. The bundles were lowered along the dorsal-ventral axis with a microdrive included as part of the bundle assembly. Single units were sorted offline using KiloSort^68^. The SC surface was identified as the depth at which visual responses were first encountered while advancing the microwire bundle. All single unit data were collected from depths within 2mm from the estimated SC surface. Only units identified at least 400um below the SC surface were included for further analysis in the current study.

Firing rates of individual neurons were represented as peristimulus time histograms (PSTHs), using 20 ms non-overlapping bins aligned to the onset of task epochs. Normalization of spike rates for each neuron was done by subtracting the mean spike count from the PSTH and dividing by the standard deviation, using the mean and standard deviation of spike counts calculated from 20 ms bins across the entire recording session. The calculation of interneuronal spike-count correlations followed previously described procedures^23^. Briefly, correlations were measured from simultaneously isolated pairs of single units with mean spike rates of at least 5 spikes/second. The Pearson correlation was computed for spike counts across trials measured during the final 200 ms of the 2-patch epoch (“delay period”).

### Mapping of visual receptive fields

SC units were further sub-selected based on having visual receptive fields that overlapped with the Gabor patch. Each visual attention task session was followed by a receptive field mapping session, during which white circular disks (118.5 cd/m^2^) of 10° in diameter were flashed against a gray background (7.2 cd/m^2^) in the visual display contralateral to the recording side. We sampled visual locations pseudo-randomly drawn from a 3 × 7 isotropic grid that extended from −25° to 25° in elevation and 0° to 90° in azimuth of the contralateral visual field. An individual trial consisted of 8 consecutive 250 ms flashes, with each flash followed by a 250 ms blank period. At least 15 flash repetitions at each grid location were presented in each mapping session.

Receptive fields of individual neurons were estimated from the mean spike counts 50-ms to 150-ms after flash onset in each grid location, after subtracting baseline activity. The baseline in each trial was defined as the mean spike count within the 100-ms period before the presentation of the first flash. Baseline-subtracted mean spike counts were normalized by dividing by the maximum value evoked across grid locations. Normalized counts were linearly interpolated between grid locations with 1° resolution using the *scatteredInterpolant* function in Matlab and smoothed with a 2D Gaussian kernel (σ_x_ = σ_y_ = 5°); we defined the receptive field boundary as the isocline at 50% of the maximum value of the smoothed, interpolated normalized counts. The area of intersection (*S_i_*) between the receptive field (*S_r_*) and the Gabor patch (*S_g_*) was used to calculate an overlap ratio (*R*), defined as *R* = ((*S_i_* / *S_r_*)^2^ + (*S_i_* / *S_g_*)^2^)^1/2^. Only units with *R* > 0.25 were used for further analysis.

### Monitoring mouse eye movements and pupil size

A high speed, 240 Hz CCD camera (ISCAN, MA) was used to monitor eye position and pupil size of head-fixed mice during the entirety of the electrophysiology experiments. We imaged an area of 1.5 mm x 3 mm with a macro lens (ISCAN, MA) centered on the eye. Four infrared light-emitting-diodes (wavelength 940 nm) were used to illuminate the eye. Commercially available acquisition software (ETL-200, ISCAN) was used to determine the center and boundary of the pupil. Eye position was obtained by subtracting the center of corneal reflection from the pupil center to compensate any translational movement of the eye in the imaging plane. The pupil displacement in 2-D image was converted to a rotation angle based on estimated eyeball radius (1.25mm) from model C57bl/6 mice^69^.

### Experimental design and statistical analysis

Data were obtained from a total number of four C57BL/6J mice in the study, two were male and two were females. We did not observe any systematic difference in behavioral performance between genders in this study.

To verify the behavioral cueing effect, we tabulated hit and false alarm rates based on the definitions of trial outcomes described for the behavioral tasks, separately for each behavioral session. Performance was then characterized by measuring sensitivity (d’) and criterion using methods from signal detection theory^70^, as follows: d’ = Φ^−1^ (H) – Φ^−1^ (F), criterion = - (Φ^−1^ (H) + Φ^−1^ (F))/2, where Φ^−1^ is the inverse of the Gaussian cumulative distribution function, H is the hit rate and F is the false alarm rate. The 95% confidence intervals (CIs) of hit and false alarm rates were computed with the *binofit* function in Matlab, which uses the Clopper-Pearson method. The 95% CIs on d’ and criterion were computed with bootstrapped resampling.

The time course of each neuron’s cue-related modulation was computed from spike counts in non-overlapping 20 ms bins (aligned on specific epochs). For each unit, the area under the receiver operating characteristic curve (auROC) was calculated between spike counts in each bin for “contra-cue” trials and counts for “ipsi-cue” trials, following methods described previously^6^.

For the cue-related modulation during the “delay-period”, spike counts from a 200-ms bin aligned on the onset of change epoch from each trial were used in two different ways. First, we computed auROC values for each neuron as described above. A bootstrapping procedure was used to compute the 95% CIs of the delay-period auROC, and if the CI was completely above or below 0.5, the unit was considered significantly modulated. Second, we computed an attention modulation index (AMI) for each unit from mean spike counts in contra-cue trials (*Count_contra_*) and ipsi-cue trials (*Count_ipsi_*) in the 200 ms epoch: AMI = (*Count_contra_* – *Count_ipsi_*) / (*Count_contra_* + *Count_ipsi_*). Nonparametric rank sum tests were performed for each unit to compare spike counts in each 200 ms bin between contra-cue and ipsi-cue trials; a unit with p < 0.05 was considered to have a significant AMI.

For the visual change-related activity, we performed the same analyses on the spike counts in the interval 60 - 210 ms after the change in orientation of the Gabor patch, computing auROC values for each neuron by comparing spike counts across trial conditions indicated in the main text (Fig. 4). Confidence intervals and associated statistical significance were again determined using a bootstrapping procedure.

For the linear regression analysis of factors contributing to spike rate variability (Fig. S2), we used Matlab function *fitlm* to model the observed spike rates in each neuron using three predictors: cue location, running speed and pupil size. We discretized running speed and pupil size, making them categorical variables. Normalized spike rate (z-scored) in four 200 ms intervals were modeled. Cue: from cue onset; early 2-patch: from 2-patch onset; mid 2-patch: from 250 ms after; delay: final 200 ms of 2-patch. Coefficients from the model fits represent the weights from each predictor that best explained the spike rate variability. The p value for testing the null hypothesis whether a predictor’s coefficient was equal to zero came from *t-statistic* for the model fit of individual unit, the degrees of freedom depended on trial counts from each recording session, between 300-800. *Post hoc* multiple comparisons with Tukey-Kramer correction after one-way ANOVA (degrees of freedom for groups: 2; degrees of freedom for errors, 930; F = 27.07) were used to assess the differences of predictor coefficient values during the delay period.

Statistical analyses were conducted in Matlab using the statistics and machine learning toolbox, and statistical significance was defined as p < 0.05 unless otherwise noted. Nonparametric rank-sum tests were computed using spike counts from −200ms to 0 ms before cue epoch onset between contralateral and ipsilateral trials to determine whether a unit display significant spatial modulation before the presence of the spatial cue. We used spike counts from four different time windows in the detection task to determine whether a unit had significant response to the onset of visual epochs: base (−100 ms to 0 ms from cue onset), cue (+50 ms to +150 ms from cue onset), late-2 (−100 ms to 0 ms from change onset), change (+50 ms to +150 ms from change onset). Nonparametric rank-sum tests were computed using spike rates in base and cue windows to determine whether a unit had a significant response to the cue. Nonparametric rank-sum tests were computed using spike rates in the late-2 and change windows to determine whether a unit had significant responses to the visual change. One-tailed nonparametric Wilcoxon signed-rank tests were performed to determine whether the distributions of population auROC values had medians significantly larger than 0.5, and whether the distributions of population AMI had medians significantly larger than 0. Paired-sample nonparametric Wilcoxon signed-rank tests were performed to compare the effect of spatial cueing on behavioral d’ and criterion across sessions. χ-square tests were performed to compare proportions of units with significant change-related auROC values across different cueing conditions. Paired-sample nonparametric Wilcoxon signed-rank tests were performed to compare the effect of spatial-cue locations on interneuronal spike count correlations across the population. The value of n reported in the figures and results indicates the number of units or unit-pairs. Error bars in figures indicate 95% CI on the median or mean, unless indicated otherwise.

## Data Availability

All of the data were acquired and initially processed using custom scripts written in Matlab (The Mathworks, MA). The Matlab code and datasets that support the findings of this study will be made available from the corresponding author upon reasonable request.

---

**Fig. S1.**
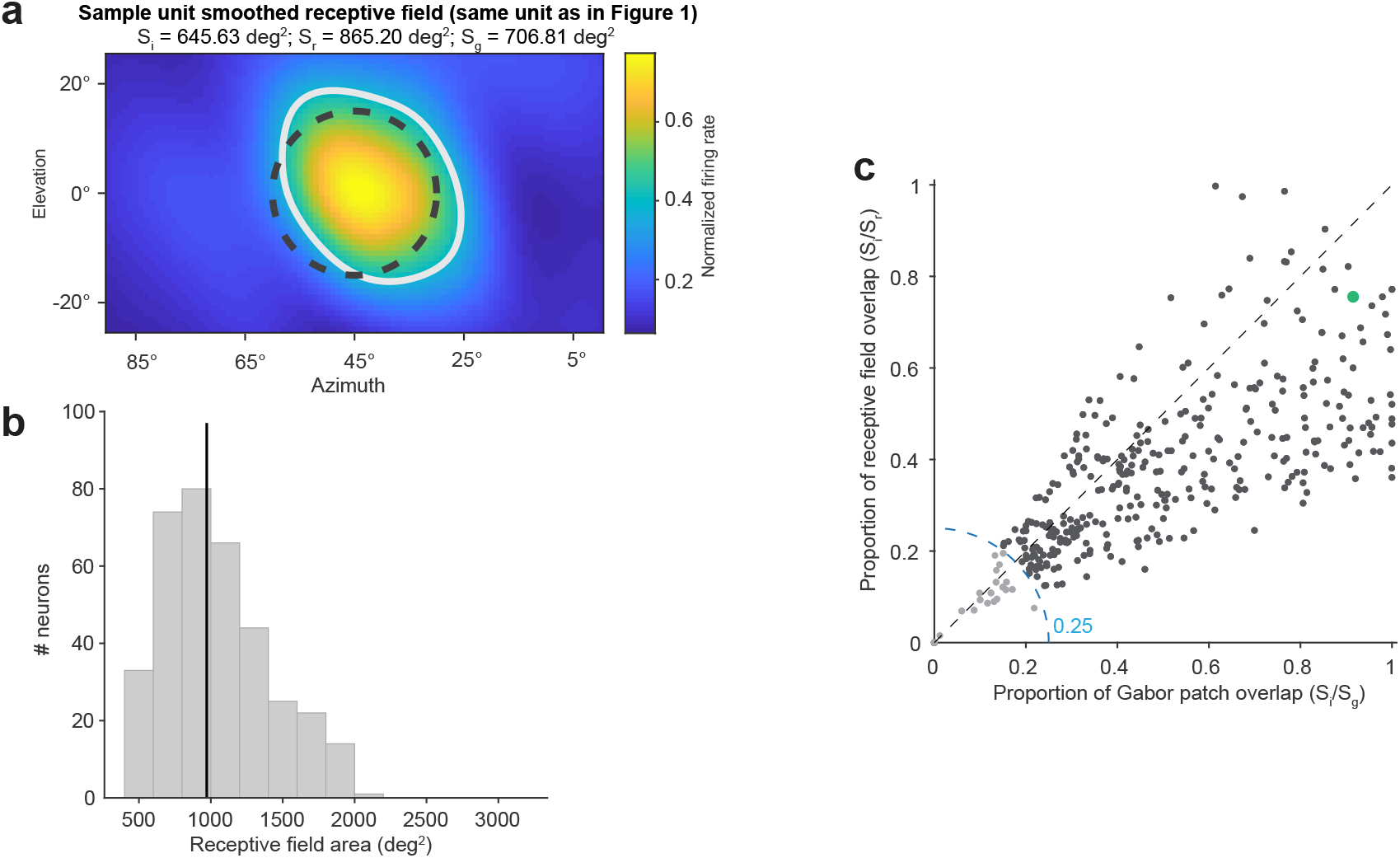
Receptive fields of recorded SC neurons. **a**) Receptive field of sample neuron shown in Fig. 1**b**. Colormap: normalized activity evoked by flashed disks across spatial grids after smoothing with 2D Gaussian kernel (***σ*** = 5°). Light gray contour line: area with at least 50% of peak activity in the smoothed map; black dashed circle: the location of Gabor patch. **b**) Distribution of receptive field size of mapped SC neurons, defined as area within the 50% contour line. Solid line: population median (971.4 deg^2^). **c**) Scatter plot of receptive field and Gabor patch overlap ratio of individual neurons. X-axis: ratio of overlapped area divided by Gabor patch area; y-axis: ratio of overlapped area divided by receptive field area. Blue dash: radius of 0.25 overlap ratio (R) used as the inclusion criteria. Only units outside the radial arc (dark dots) were used for further analysis in the paper. The larger green dot is the same unit shown in panel **a**.

**Fig. S2.**
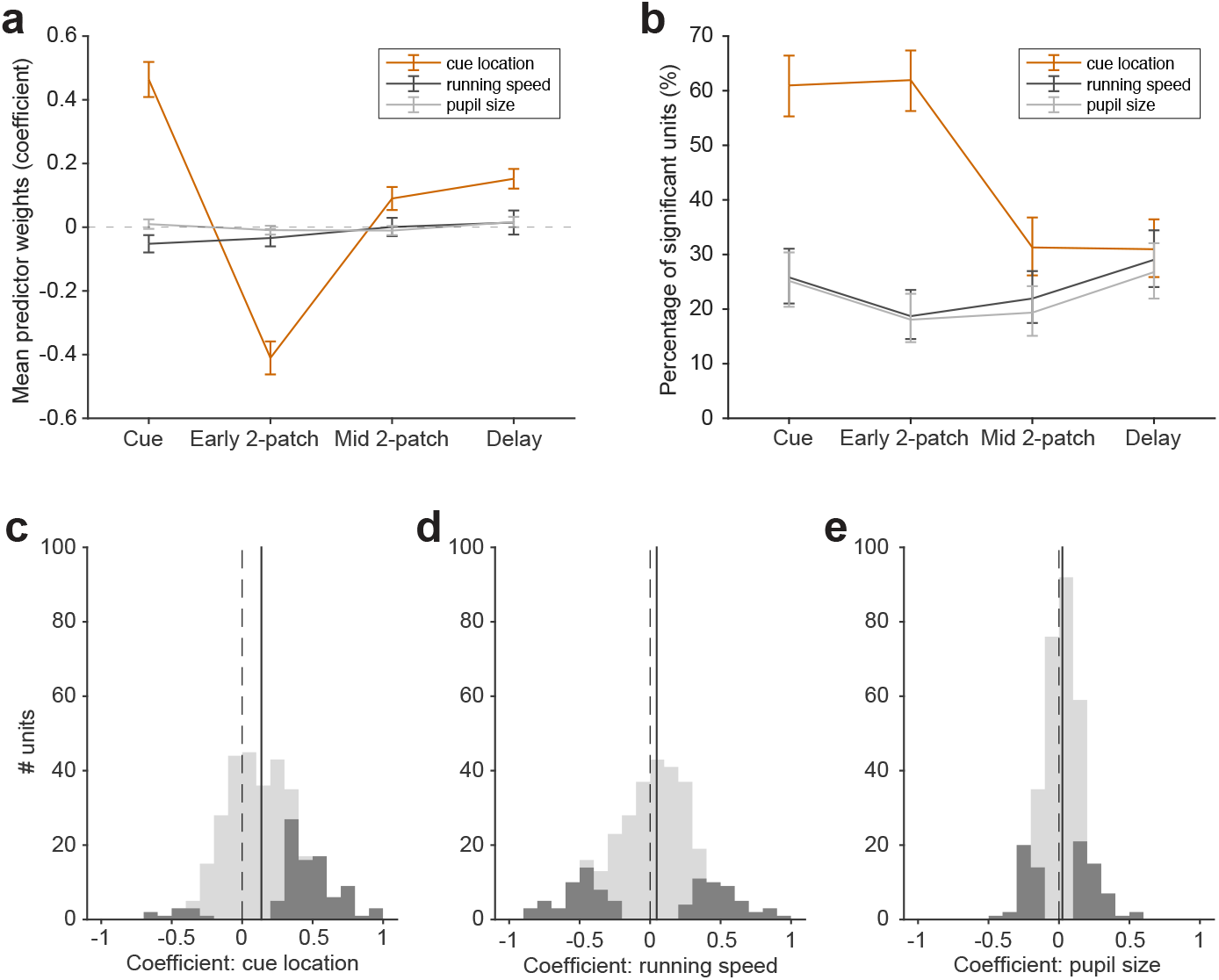
Contribution of running speed, pupil size and cue location on SC activity during the task. **a**) Mean coefficient of linear regression of cue location (brown), running speed (black) and pupil size (gray) on variability of SC activity during different 200 ms epoch intervals. Cue: from cue onset; early 2-patch: from 2-patch onset; mid 2-patch: from 250ms after; delay: last 200ms of 2-patch. Errorbar: 95% CI. **b**) Similar to **a**, but for percentage of units significantly modulated by different predictors (p < 0.01, t-statistic to test the null hypothesis whether a predictor coefficient in the regression model is equal to zero for each unit, see Methods). **c**) Distribution of coefficient of cue location on SC activity during delay period. Dark bars are units with coefficient significantly different from zero. Dashed line indicates null value of 0, solid line indicates population median (0.14). **d**) Presentation as in **c**, but for running speed (median = 0.05). **e**) Presentation as in **c**, but for pupil size (median = 0.02).

